# HiMaLAYAS: enrichment-based annotation and visualization of hierarchically clustered matrices

**DOI:** 10.64898/2026.02.11.705303

**Authors:** Ira Horecka, Hannes Röst

**Affiliations:** Terrence Donnelly Centre for Cellular & Biomolecular Research, University of Toronto, Toronto, Ontario M5S 3E1, Canada; Department of Molecular Genetics, University of Toronto, Toronto, Ontario M5S 1A8, Canada; Department of Computer Science, University of Toronto, Toronto, Ontario M5S 2E4, Canada

## Abstract

**Summary:** Hierarchical clustering reveals structure in high-dimensional biological matrices, but the resulting dendrogram-defined clusters are commonly visualized rather than tested for annotation enrichment. Existing workflows often require separate tools to cluster data, test enrichment, and display heatmaps. Here, we introduce Hierarchical Matrix Layout and Annotation Software (HiMaLAYAS), a Python framework for post hoc enrichment-based annotation and visualization of hierarchically clustered matrices. HiMaLAYAS treats dendrogram-defined clusters as statistical units, tests categorical annotations for enrichment, controls multiple testing, and renders significant annotations alongside clusters in a single workflow. Applied to a *Saccharomyces cerevisiae* genetic interaction profile similarity matrix, HiMaLAYAS annotated biological-process enrichment at parent-cluster and zoomed-cluster levels. Parent-level annotations persisted across clustering-parameter sweeps, exceeded null expectations from randomized cluster membership and annotation labels, and remained stable under matrix perturbation. HiMaLAYAS also annotated clusters in a non-biological country-sector input-output matrix, extending its application beyond biology.

**Availability and Implementation:** HiMaLAYAS is a Python package available via pip, distributed under the BSD 3-Clause License at https://github.com/himalayas-base/himalayas, and archived on Zenodo at https://doi.org/10.5281/zenodo.18610373.

## 1 Introduction

Hierarchical clustering organizes high-dimensional biological matrices by ordering rows and columns into contiguous regions of related observations while color encodes matrix values. This enables visual interpretation across thousands of measurements and is widely adopted for exploratory analysis in functional genomics and systems biology (Eisen *et al*., 1998; Spellman *et al*., 1998).

Clustering creates a dendrogram that is typically cut to define clusters (Langfelder and Horvath, 2008), which are often visualized or annotated with pathway membership tracks alongside the matrix (Gu *et al*., 2016). Dendrogram-defined clusters, however, are not always tested for enrichment.

Available tools can cluster data, test enrichment, and display heatmaps, but these capabilities are often distributed across separate packages rather than provided as a single workflow. Because enrichment annotations depend on cluster definition, users also need to know whether annotations persist across clustering choices, exceed null expectations from randomized cluster membership and annotation labels, and remain stable under matrix perturbation.

We introduce Hierarchical Matrix Layout and Annotation Software (HiMaLAYAS), a Python framework for post hoc enrichment-based annotation and visualization of hierarchically clustered matrices across biological and non-biological domains. HiMaLAYAS treats dendrogram-defined clusters as statistical units, tests categorical annotations for enrichment, controls multiple testing, and renders significant annotations alongside clusters.

We apply HiMaLAYAS to a *Saccharomyces cerevisiae* genetic interaction profile similarity matrix from Costanzo *et al*. (2016) and show enriched annotations at parent-cluster and zoomed-cluster levels. We evaluate robustness using clustering-parameter sweeps, null models, and matrix-noise perturbation; compare the workflow with related tools; and demonstrate portability using a non-biological country-sector input-output matrix.

## 2 Methods

### 2.1 Input matrices and cluster definition

HiMaLAYAS hierarchically clusters a real-valued matrix capturing relationships among observations (Fig. 1A). We expect users to preprocess the input matrix to a scale appropriate for hierarchical clustering, such as a similarity matrix, normalized expression matrix, or transformed count-like matrix. Users can configure linkage method, distance metric, and dendrogram distance threshold in HiMaLAYAS. After clustering, users cut the dendrogram to define cluster number and membership, then specify a minimum cluster size for enrichment testing. By default, HiMaLAYAS merges clusters below this size upward along the dendrogram until the minimum size is met; alternatively, it can preserve the hierarchy but exclude these clusters from enrichment testing.

**Figure 1.**
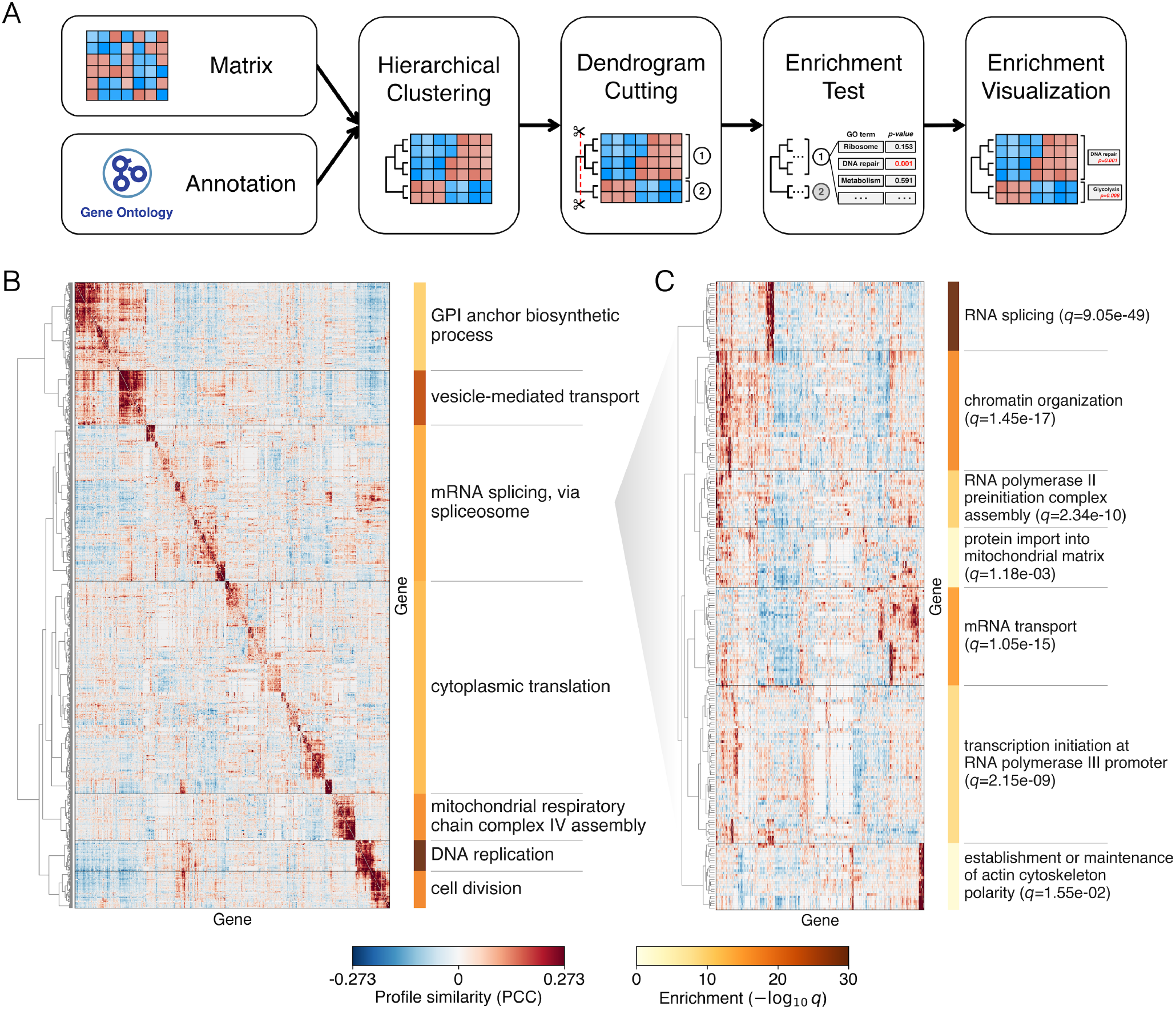
HiMaLAYAS workflow and its application to a *Saccharomyces cerevisiae* genetic interaction profile similarity matrix (Costanzo *et al*., 2016). (**A**) HiMaLAYAS workflow. A real-valued matrix and categorical annotations serve as inputs. HiMaLAYAS hierarchically clusters the matrix, cuts the dendrogram at a user-defined distance threshold, tests categorical annotations for enrichment, controls multiple testing, and renders significant annotations alongside clusters. (**B**) HiMaLAYAS applied to the yeast genetic interaction profile similarity matrix, focusing on 1,053 genes with high profile variance. Each matrix entry represents two genes’ similarity in their genetic interaction profiles measured by Pearson correlation. Dendrogram-defined clusters were tested for Gene Ontology Biological Process (GO BP; Ashburner *et al*., 2000) enrichment, with top-ranked significant annotations rendered alongside clusters. (**C**) HiMaLAYAS applied to a zoomed view of the parent cluster enriched for mRNA splicing, resolving finer clusters within the parent cluster. Dendrogram-defined clusters in the zoomed view were tested for GO BP enrichment, with top-ranked significant annotations rendered alongside clusters.

### 2.2 Enrichment testing and visualization

HiMaLAYAS tests enrichment for categorical annotations across clusters. Annotations define term-to-observation memberships, such as Gene Ontology Biological Process (GO BP; Ashburner *et al*., 2000) terms associated with genes. HiMaLAYAS defines the enrichment background from matrix observations and retains this background when a selected cluster is reanalyzed in a zoomed view. Each observation can belong to multiple annotation terms, and HiMaLAYAS tests only cluster–term pairs meeting a user-defined minimum member overlap. It applies a one-sided hypergeometric test to determine whether a term’s members are overrepresented among a cluster’s observations. HiMaLAYAS controls multiple testing across cluster–term tests using the Benjamini–Hochberg false discovery rate (FDR) procedure (Benjamini and Hochberg, 1995). In this study, we consider cluster–term enrichments significant at FDR ≤ 0.05, and we rank significant annotations by q-value.

HiMaLAYAS renders the top-ranked significant annotation alongside each cluster but retains the full cluster– term result set. HiMaLAYAS does not, however, filter semantically redundant Gene Ontology terms before visualization; users can group such terms with tools such as simplifyEnrichment (Gu and Hübschmann, 2023). HiMaLAYAS supports annotated matrix views, condensed hierarchy views, post hoc row data tracks, and configurable text styling and figure layout. It exports figures in raster or vector formats for publication.

### 2.3 Robustness and null-analysis design

Because HiMaLAYAS tests enrichment on dendrogram-defined clusters, we tested whether recovered annotations varied with clustering choices, null structure, or matrix perturbation. We used the *Saccharomyces cerevisiae* genetic interaction profile similarity matrix (Costanzo *et al*., 2016) shown in Fig. 1B as the reference analysis. In the reference analysis, we used a dendrogram distance threshold of 16 and a minimum cluster size of 30.

To test sensitivity to clustering choices, we varied the dendrogram distance threshold and minimum cluster size, reran enrichment testing, and scored recovery of each of the seven reference headline GO BP terms using the cluster with the greatest gene overlap with its reference cluster.

To test whether the observed enrichment signal exceeded null expectation, we used two 1,000-replicate null models: same-size random clusters with annotations fixed and annotation-label permutations with clusters fixed. Each replicate used the same enrichment test and FDR correction as the reference analysis.

Finally, to test sensitivity to matrix perturbation, we added symmetric zero-mean Gaussian noise at fractions of 0.05, 0.10, 0.20, 0.30, and 0.50 of the off-diagonal matrix standard deviation, reclustered the perturbed matrix, and reran enrichment testing. We scored recovery of each of the seven reference headline GO BP terms using the perturbed cluster with the greatest gene overlap with its reference cluster.

## 3 Results

### 3.1 Annotating the *Saccharomyces cerevisiae* genetic interaction profile similarity matrix

We applied HiMaLAYAS to a *Saccharomyces cerevisiae* genetic interaction profile similarity matrix from Costanzo *et al*. (2016), focusing on 1,053 genes with high profile variance (Fig. 1B). Dendrogram-defined clusters significantly enriched for Gene Ontology Biological Process (GO BP; Ashburner *et al*., 2000) terms were annotated alongside each cluster. We identified parent clusters enriched for vesicle-mediated transport, mRNA splicing, and DNA replication. We then applied HiMaLAYAS to a zoomed view of the parent cluster enriched for mRNA splicing, identifying finer clusters enriched for mRNA transport, RNA splicing, and chromatin organization (Fig. 1C). HiMaLAYAS thereby resolves a broad annotated parent cluster into finer functional subclusters while retaining the same enrichment background.

Because dense matrices can obscure dendrogram structure, HiMaLAYAS summarizes annotated clusters with condensed hierarchy views and links cluster annotations to row-level data through post hoc row data tracks, as shown in the parent-level and zoomed yeast analyses (Supplementary Fig. S1).

### 3.2 Testing annotation robustness

Clustering determines the groups tested for enrichment. We tested annotation sensitivity to dendrogram distance-threshold and minimum-cluster-size parameters using the seven headline GO BP terms from the parent-level yeast analysis in Fig. 1B. These terms were largely recovered across parameter sweeps, indicating that the headline annotations were not dependent on a single dendrogram cut or minimum-size setting (Fig. 2A,B).

**Figure 2.**
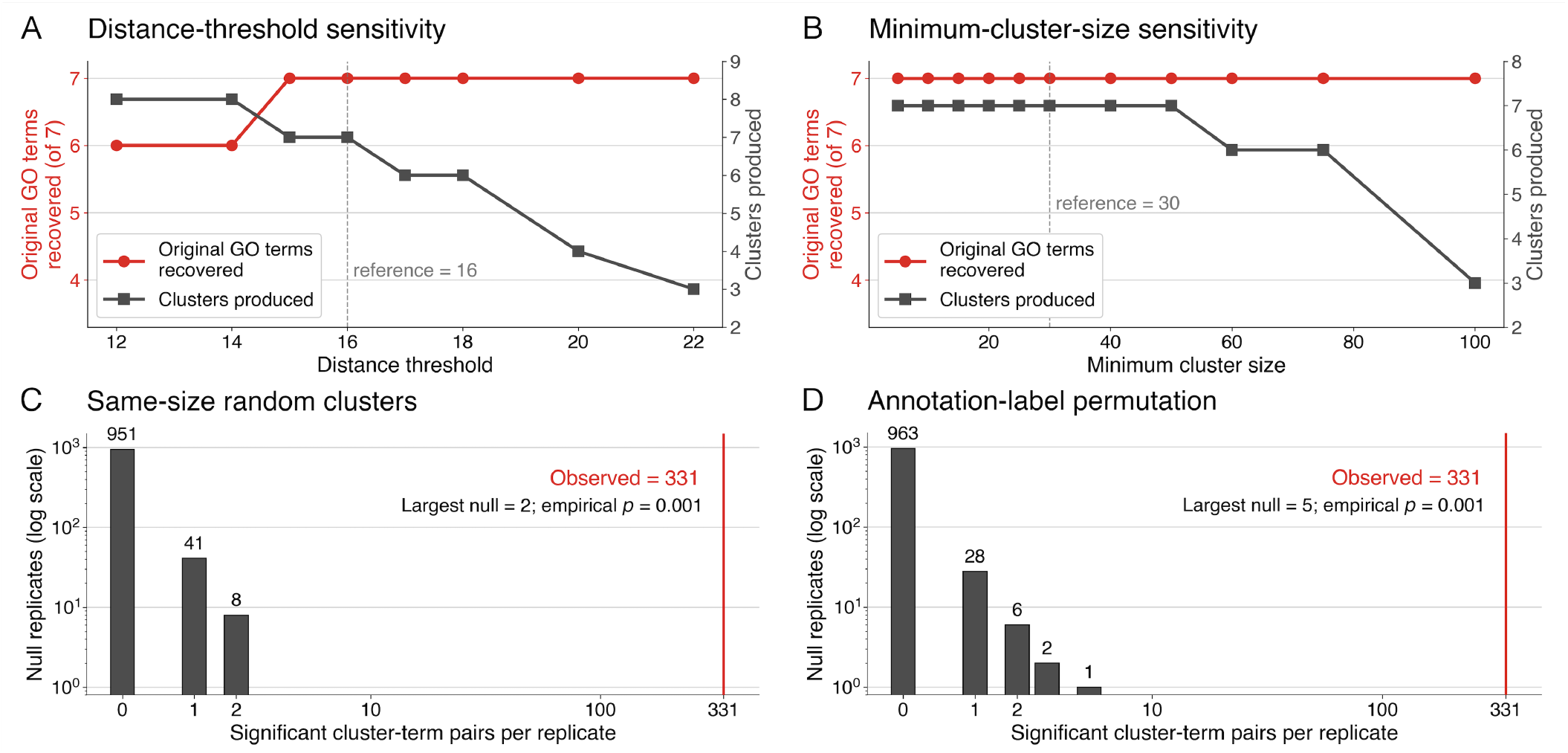
Robustness and null analysis of Gene Ontology Biological Process (GO BP; Ashburner *et al*., 2000) annotations in the *Saccharomyces cerevisiae* genetic interaction profile similarity matrix (Costanzo *et al*., 2016). (**A**,**B**) Sensitivity of the Fig. 1B reference analysis to dendrogram distance threshold (**A**) and minimum cluster size (**B**). Red curves show recovery of the seven reference headline GO BP terms after rerunning enrichment testing. Gray curves show the number of clusters produced at each setting, and dashed lines mark the reference settings. A headline term was counted as recovered when it remained significant in the cluster with the greatest gene overlap with its reference cluster. (**C**,**D**) Null analysis of the observed enrichment signal. The reference analysis produced 331 significant cluster–term pairs (q ≤ 0.05), shown as a red vertical line. The 1,000-replicate null distributions were generated using same-size random clusters with annotations fixed (**C**) and annotation-label permutations with clusters fixed (**D**). Each replicate used the same enrichment test and FDR correction as the reference analysis.

Significant annotations can arise from arbitrary cluster membership or annotation labels. We tested the parent-level yeast analysis against two 1,000-replicate null models: randomized cluster membership and randomized annotation labels. The parent-level yeast analysis produced 331 significant cluster–term pairs, exceeding the largest null value by more than 60-fold (5; empirical *P* = 0.001 for both; Fig. 2C,D).

Finally, matrix noise can alter cluster composition. We tested whether the headline annotations were recovered after perturbing the parent-level yeast matrix across five noise fractions and five replicates per fraction. The headline terms ranked first in 138 of 175 perturbed analyses and remained significant in 174 of 175 analyses (Supplementary Fig. S2).

### 3.3 A single workflow for annotating and visualizing hierarchically clustered matrices

Related tools such as WGCNA (Langfelder and Horvath, 2008), clusterProfiler (Yu *et al*., 2012), ComplexHeatmap (Gu *et al*., 2016), VisHiC (Krushevskaya *et al*., 2009), funcExplorer (Kolberg *et al*., 2018), and Clustergrammer (Fernandez *et al*., 2017) collectively support clustering, enrichment testing, and heatmap visualization. These capabilities, however, are often distributed across multiple tools. HiMaLAYAS packages dendrogram cutting, enrichment testing, multiple-testing control, and annotation visualization into a single workflow (Supplementary Fig. S3).

Dense matrices with categorical metadata appear beyond biology. To test this portability, we applied HiMaLAYAS to the non-biological 2014 World Input-Output Database country-sector matrix (Timmer *et al*., 2015). HiMaLAYAS clustered origin country-sectors by their intermediate-flow profiles and identified clusters significantly enriched for shared origin countries (Supplementary Fig. S4).

## 4 Conclusion

HiMaLAYAS provides a single workflow for enrichment-based annotation and visualization of hierarchically clustered matrices. In the yeast genetic interaction profile similarity matrix, HiMaLAYAS annotated dendrogram-defined clusters enriched for biological processes at parent-cluster and zoomed-cluster levels. In the parent-level yeast analysis, headline annotations were robust to clustering-parameter changes and matrix noise and exceeded null expectations from randomized cluster membership and annotation labels. Finally, we applied HiMaLAYAS to a non-biological matrix, extending the workflow beyond biology.

## Supplementary data

Supplementary data are available at *Bioinformatics Advances* online.

## Data availability

The HiMaLAYAS software is available at https://github.com/himalayas-base/himalayas. Documentation and Jupyter notebook tutorials are available at https://github.com/himalayas-base/himalayas-docs. The datasets and workflows used to generate the figures in this study are available at https://github.com/himalayas-base/himalayas-publication.

## References

Ashburner M, Ball CA, Blake JA, et al. Gene Ontology: tool for the unification of biology. Nat Genet 2000;25:25–29.

Benjamini Y, Hochberg Y. Controlling the false discovery rate: a practical and powerful approach to multiple testing. J R Stat Soc Series B 1995;57:289–300.

Costanzo M, VanderSluis B, Koch EN, et al. A global genetic interaction network maps a wiring diagram of cellular function. Science 2016;353:aaf1420.

Eisen MB, Spellman PT, Brown PO, Botstein D. Cluster analysis and display of genome-wide expression patterns. Proc Natl Acad Sci 1998;95:14863–14868.

Fernandez NF, Gundersen GW, Rahman A, et al. Clustergrammer, a web-based heatmap visualization and analysis tool for high-dimensional biological data. Sci Data 2017;4:170151.

Gu Z, Eils R, Schlesner M. Complex heatmaps reveal patterns and correlations in multidimensional genomic data. Bioinformatics 2016;32:2847–2849.

Gu Z, Hübschmann D. simplifyEnrichment: a Bioconductor package for clustering and visualizing functional enrichment results. Genomics Proteomics Bioinformatics 2023;21:190–202.

Kolberg L, Kuzmin I, Adler P, Vilo J, Peterson H. funcExplorer: a tool for fast data-driven functional characterisation of high-throughput expression data. BMC Genomics 2018;19:817.

Krushevskaya D, Peterson H, Reimand J, Kull M, Vilo J. VisHiC—hierarchical functional enrichment analysis of microarray data. Nucleic Acids Res 2009;37:W587–W592.

Langfelder P, Horvath S. WGCNA: an R package for weighted correlation network analysis. BMC Bioinformatics 2008;9:559.

Spellman PT, Sherlock G, Zhang MQ, et al. Comprehensive identification of cell cycle–regulated genes of the yeast Saccharomyces cerevisiae by microarray hybridization. Mol Biol Cell 1998;9:3273–3297.

Timmer MP, Dietzenbacher E, Los B, Stehrer R, de Vries GJ. An illustrated user guide to the World Input-Output Database: the case of global automotive production. Rev Int Econ 2015;23:575–605.

Yu G, Wang LG, Han Y, He QY. clusterProfiler: an R package for comparing biological themes among gene clusters. OMICS 2012;16:284–287.

