## Supplementary Figures for "HiMaLAYAS: enrichment-based annotation and visualization of hierarchically clustered matrices"

This document contains supplementary figures presented in *HiMaLAYAS: enrichment-based annotation and visualization of hierarchically clustered matrices*.

- **Supplementary Figure S1:** Annotated matrix and condensed hierarchy views of the *Saccharomyces cerevisiae* genetic interaction profile similarity matrix.
- **Supplementary Figure S2:** Matrix-perturbation robustness of seven headline Gene Ontology Biological Process annotations in the *Saccharomyces cerevisiae* genetic interaction profile similarity matrix.
- **Supplementary Figure S3:** Related-tool capability comparison for annotating and visualizing hierarchically clustered matrices.
- **Supplementary Figure S4:** Non-biological matrix portability example using the World Input-Output Database.

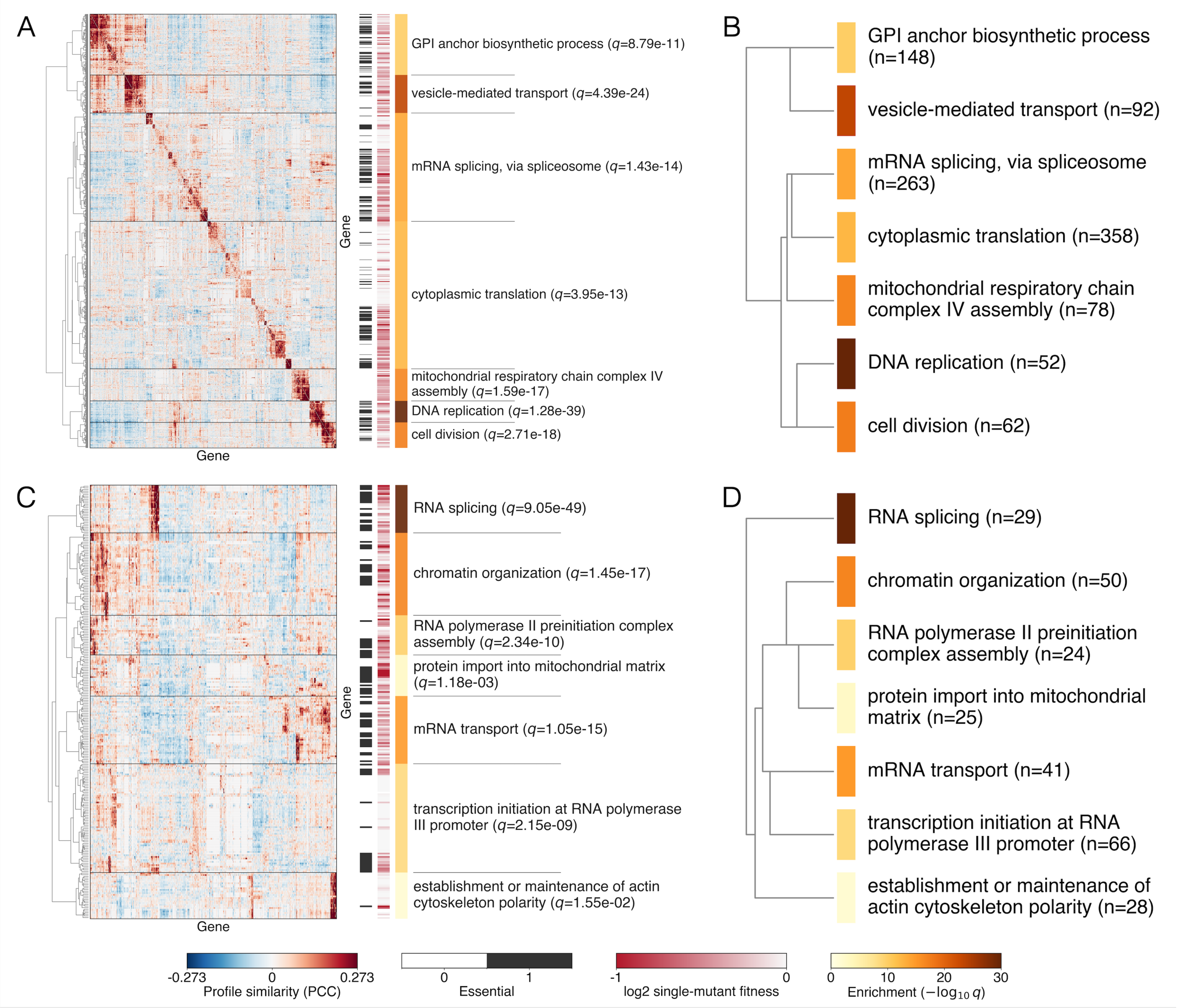


**Supplementary Figure S1.** Annotated matrix and condensed hierarchy views of the *Saccharomyces cerevisiae* genetic interaction profile similarity matrix (Costanzo *et al*., 2016). (**A,B**) HiMaLAYAS applied to the yeast genetic interaction profile similarity matrix, focusing on 1,053 genes with high profile variance, shown as an annotated matrix view (**A**) and condensed hierarchy view (**B**). (**C,D**) HiMaLAYAS applied to a zoomed view of the parent cluster enriched for mRNA splicing, shown as an annotated matrix view (**C**) and condensed hierarchy view (**D**). Annotated matrix views include post hoc row data tracks for ORF essentiality and single-mutant fitness at 30°C from Costanzo *et al.* (2016). For each labeled cluster, the top-ranked enriched Gene Ontology Biological Process (GO BP; Ashburner *et al*., 2000) term by q-value is shown. Condensed hierarchy views summarize the same dendrogram-defined clusters and significant annotations shown in the annotated matrix views.

#### Cluster 1
GPI anchor biosynthetic process ($q$=8.79e-11)

| Noise Fraction | #1 | Top-5 (not #1) | Not Significant |
|---|---|---|---|
| 0.05 | 5 | 0 | 0 |
| 0.10 | 5 | 0 | 0 |
| 0.20 | 4 | 1 | 0 |
| 0.30 | 5 | 0 | 0 |
| 0.50 | 5 | 0 | 0 |
| Pooled (25) | 24 | 1 | 0 |

#### Cluster 2
vesicle-mediated transport ($q$=4.39e-24)

| Noise Fraction | #1 | Top-5 (not #1) | Not Significant |
|---|---|---|---|
| 0.05 | 5 | 0 | 0 |
| 0.10 | 5 | 0 | 0 |
| 0.20 | 5 | 0 | 0 |
| 0.30 | 5 | 0 | 0 |
| 0.50 | 5 | 0 | 0 |
| Pooled (25) | 25 | 0 | 0 |

#### Cluster 3
mRNA splicing, via spliceosome ($q$=1.43e-14)

| Noise Fraction | #1 | Top-5 (not #1) | Not Significant |
|---|---|---|---|
| 0.05 | 5 | 0 | 0 |
| 0.10 | 3 | 2 | 0 |
| 0.20 | 4 | 1 | 0 |
| 0.30 | 2 | 3 | 0 |
| 0.50 | 4 | 1 | 0 |
| Pooled (25) | 18 | 7 | 0 |

#### Cluster 4
cytoplasmic translation ($q$=3.95e-13)

| Noise Fraction | #1 | Top-5 (not #1) | Not Significant |
|---|---|---|---|
| 0.05 | 3 | 1 | 1 |
| 0.10 | 3 | 2 | 0 |
| 0.20 | 0 | 5 | 0 |
| 0.30 | 0 | 5 | 0 |
| 0.50 | 2 | 3 | 0 |
| Pooled (25) | 8 | 16 | 1 |

#### Cluster 5
mitochondrial respiratory chain complex IV assembly ($q$=1.59e-17)

| Noise Fraction | #1 | Top-5 (not #1) | Not Significant |
|---|---|---|---|
| 0.05 | 5 | 0 | 0 |
| 0.10 | 5 | 0 | 0 |
| 0.20 | 5 | 0 | 0 |
| 0.30 | 5 | 0 | 0 |
| 0.50 | 5 | 0 | 0 |
| Pooled (25) | 25 | 0 | 0 |

#### Cluster 6
DNA replication ($q$=1.28e-39)

| Noise Fraction | #1 | Top-5 (not #1) | Not Significant |
|---|---|---|---|
| 0.05 | 5 | 0 | 0 |
| 0.10 | 5 | 0 | 0 |
| 0.20 | 5 | 0 | 0 |
| 0.30 | 5 | 0 | 0 |
| 0.50 | 5 | 0 | 0 |
| Pooled (25) | 25 | 0 | 0 |

#### Cluster 7
cell division ($q$=2.71e-18)

| Noise Fraction | #1 | Top-5 (not #1) | Not Significant |
|---|---|---|---|
| 0.05 | 4 | 1 | 0 |
| 0.10 | 4 | 1 | 0 |
| 0.20 | 2 | 3 | 0 |
| 0.30 | 3 | 2 | 0 |
| 0.50 | 0 | 5 | 0 |
| Pooled (25) | 13 | 12 | 0 |

**Supplementary Figure S2.** Matrix-perturbation robustness of seven headline Gene Ontology Biological Process (GO BP; Ashburner *et al.*, 2000) annotations in the *Saccharomyces cerevisiae* genetic interaction profile similarity matrix (Costanzo *et al.*, 2016). Matrix perturbation was implemented by adding symmetric zero-mean Gaussian noise to the yeast matrix at fractions of 0.05, 0.10, 0.20, 0.30, and 0.50 of the off-diagonal matrix standard deviation; diagonal values were unchanged. For each of the seven reference clusters from the parent-level yeast analysis, we tracked the original headline GO BP term after matrix perturbation, reclustering, and enrichment testing. We scored recovery using the perturbed cluster with the greatest gene overlap with its reference cluster. Each table summarizes one reference cluster and reports whether the original headline term remained the top-ranked significant term by q-value, remained significant within the top five but was not top-ranked, or was not significant. Each noise fraction includes five replicate analyses, and the pooled row summarizes all 25 replicates for that cluster. The original headline terms ranked first in 138 of 175 perturbed analyses and remained significant in 174 of 175 analyses.

| | Hierarchical zoom | Condensed dendrogram | Beyond expression data | Enrichment test | Multiple-testing control | Post hoc annotation | Local install / maintained |
|---|---|---|---|---|---|---|---|
| HiMaLAYAS | ● | ● | ● | ● | ● | ● | ● |
| funcExplorer | ● | ● | ● | ● | ● | ○ | ◑ |
| Clustergrammer | ● | ● | ● | ◑ | ○† | ◑ | ◑ |
| VisHiC | ● | ● | ○ | ● | ● | ○ | ○ |
| Heatmap + manual ORA | ◑ | ○ | ◑ | ● | ● | ◑ | ○ |
| clusterProfiler | ○ | ○ | ○ | ● | ● | ◑ | ● |
| WGCNA | ○ | ○ | ◑ | ◑ | ◑ | ◑ | ● |
| ComplexHeatmap | ○ | ○ | ● | ○ | ○ | ○ | ● |

● Yes
◑ Partial
○ No
○ Not documented (†)

**Supplementary Figure S3.** HiMaLAYAS compared with related tools and workflows for annotating and visualizing hierarchically clustered matrices. The qualitative comparison focuses on capabilities relevant to clustering, enrichment testing, and heatmap visualization. Columns summarize seven capabilities: hierarchical zoom, condensed dendrogram views, support beyond expression data, enrichment testing, multiple-testing control, post hoc annotation, and local installation or maintenance. Rows compare HiMaLAYAS, funcExplorer (Kolberg *et al.*, 2018), Clustergrammer (Fernandez *et al.*, 2017), VisHiC (Krushevskaya *et al.*, 2009), Heatmap + manual ORA, clusterProfiler (Yu *et al.*, 2012), WGCNA (Langfelder and Horvath, 2008), and ComplexHeatmap (Gu *et al.*, 2016). Filled circles indicate *Yes*, half-filled circles indicate *Partial*, open circles indicate *No*, and open circles with a dagger indicate *Not documented*. Classifications reflect documented support for the workflow assessed here and provide qualitative positioning rather than a runtime, accuracy, biological-discovery, or usability benchmark.

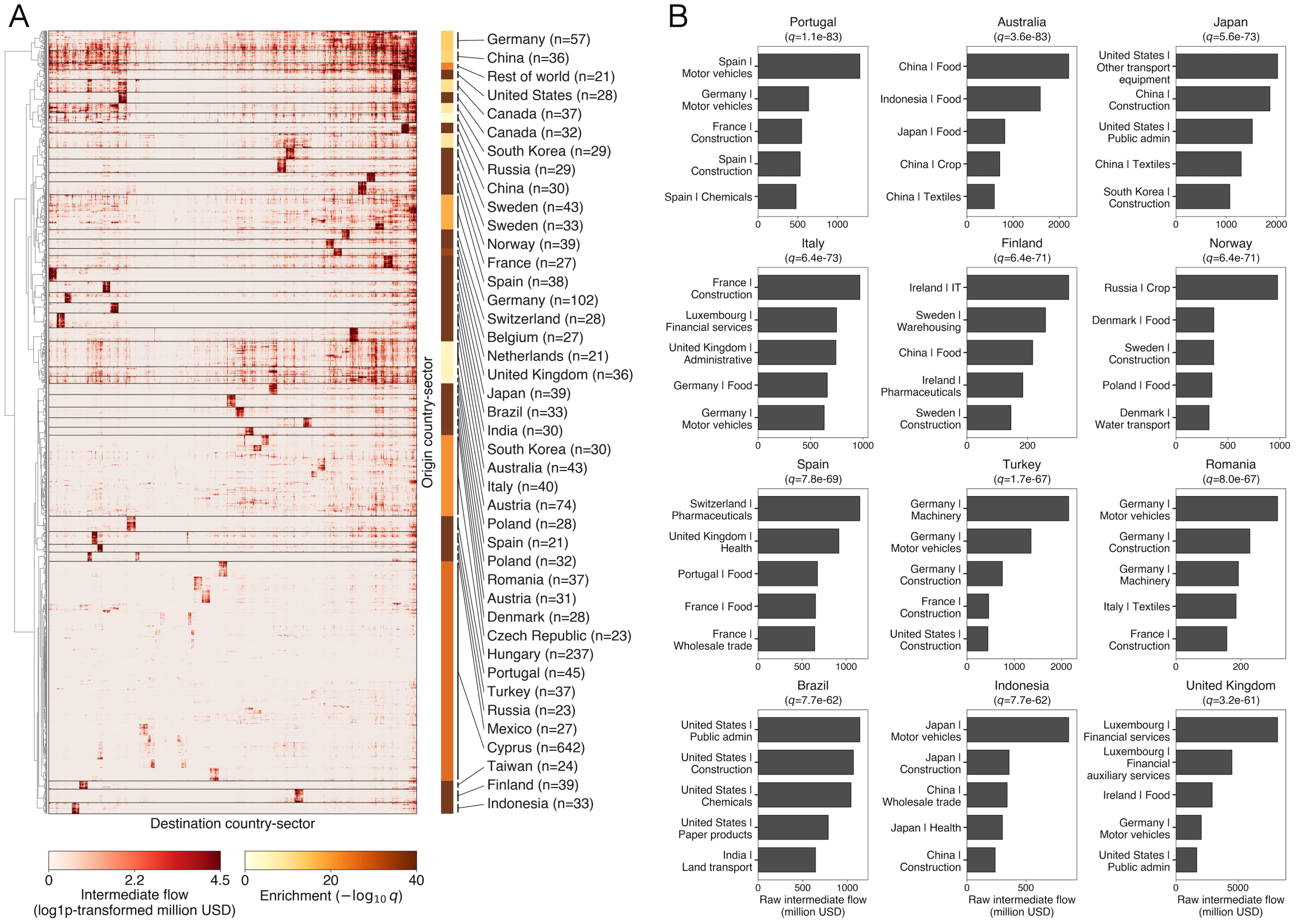


**Supplementary Figure S4.** Non-biological matrix portability example using the 2014 World Input-Output Database (WIOD; Timmer *et al*., 2015) country-sector input-output matrix. (**A**) Annotated matrix view. Rows represent origin country-sectors, columns represent destination country-sectors, and matrix values represent log1p-transformed intermediate flows in million USD. HiMaLAYAS clustered origin country-sectors by their intermediate-flow profiles and tested row clusters for shared origin-country enrichment, with top-ranked significant annotations rendered alongside clusters. (**B**) Descriptive summaries of selected origin-country-enriched clusters. For each cluster, bars show the destination country-sector columns with the largest summed cross-border intermediate flows across member origin country-sector rows.

### References

Ashburner M, Ball CA, Blake JA, *et al.* Gene Ontology: tool for the unification of biology. *Nat Genet* 2000;**25**:25–29.

Costanzo M, VanderSluis B, Koch EN, *et al.* A global genetic interaction network maps a wiring diagram of cellular function. *Science* 2016;**353**:aaf1420.

Fernandez NF, Gundersen GW, Rahman A, *et al.* Clustergrammer, a web-based heatmap visualization and analysis tool for high-dimensional biological data. *Sci Data* 2017;**4**:170151.

Gu Z, Eils R, Schlesner M. Complex heatmaps reveal patterns and correlations in multidimensional genomic data. *Bioinformatics* 2016;**32**:2847–2849.

Kolberg L, Kuzmin I, Adler P, Vilo J, Peterson H. funcExplorer: a tool for fast data-driven functional characterisation of high-throughput expression data. *BMC Genomics* 2018;**19**:817.

Krushevskaya D, Peterson H, Reimand J, Kull M, Vilo J. VisHiC—hierarchical functional enrichment analysis of microarray data. *Nucleic Acids Res* 2009;**37**:W587–W592.

Langfelder P, Horvath S. WGCNA: an R package for weighted correlation network analysis. *BMC Bioinformatics* 2008;**9**:559.

Timmer MP, Dietzenbacher E, Los B, Stehrer R, de Vries GJ. An illustrated user guide to the World Input-Output Database: the case of global automotive production. *Rev Int Econ* 2015;**23**:575–605.

Yu G, Wang LG, Han Y, He QY. clusterProfiler: an R package for comparing biological themes among gene clusters. *OMICS* 2012;**16**:284–287.
